# Phase separation in amino acid mixtures is governed by composition

**DOI:** 10.1101/2022.03.02.482633

**Authors:** David De Sancho

## Abstract

Macromolecular phase separation has recently come to immense prominence as it is central to the formation of membraneless organelles, leading to a new paradigm of cellular organization. This type of phase transition, often termed liquid-liquid phase separation (LLPS), is mediated by molecular interactions between biomolecules, including nucleic acids and both ordered and disordered proteins. In the latter case, the separation between protein-dense and dilute phases is often interpreted using models adapted from polymer theory. Specifically, the “stickers and spacers” model proposes that the formation of condensate-spanning networks in protein solutions originates from the interplay between two classes of residues and that the main determinants for phase separation are multivalency and sequence patterning. The duality of roles of stickers (aromatics like Phe and Tyr) and spacers (Gly and polar residues) may apply more broadly in protein-like mixtures, and the presence of these two types of components alone may suffice for LLPS to take place. In order to explore this hypothesis, we use atomistic molecular dynamics simulations of capped amino-acid residues as a minimal model system. We study the behaviour of pure amino acids in water for three types of residues corresponding to the spacer and sticker categories, and their multicomponent mixtures. In agreement with previous observations, we find that the spacer-type amino acids fail to phase-separate on their own, while the sticker is prone to aggregation. However, ternary amino acid mixtures involving both types of amino acids phase-separate into two phases that retain intermediate degrees of compaction and greater fluidity than sticker-only condensates. Our results suggest that LLPS is an emergent property of amino acid mixtures determined by composition.

## Introduction

Cellular organisms are well known to use lipid membranes to compartmentalise their many functions into organelles. In the last decade, it has become apparent that organelles lacking membranes are also ubiquitous in cells.^1^ These membraneless organelles include the nucleolus, stress granules, nuclear speckles and Cajal bodies, among many others. They are rich in proteins and also nucleic acids,^2^ which form molecular condensates as they undergo a type of phase transition usually called liquid-liquid phase separation (LLPS). This term is used because, as a result, a dilute and a concentrated phase are formed, both of which have liquid-like properties such as fusion into larger droplets, flow in response to stress, or wetting and dropping behaviours typical of liquids.^3^ In fact, liquid-liquid phase separation may be more accurately described as the formation of condensate-spanning networks, due to the interplay between the propensity of the biomolecule to both phase-separate and percolate.^4,5^ When condensates are formed following phase separation coupled to percolation (PSCP), they are viscoelastic in nature rather than simple liquids.^6^ The arrival to prominence of membraneless organelles has sparked considerable interest in the types of molecules undergoing LLPS and the physical principles governing this process.^7^ Although this type of phase transition has been reported for modular folded domains,^8^ many proteins that phase separate are intrinsically disordered or have intrinsically disordered regions. These IDPs or IDRs have strong biases in sequence composition,^9–11^ as in the case of the “low complexity regions” (LCRs) that are abundant in biomolecular condensates.^12^

Because polypeptide chains are often the drivers of LLPS, concepts from polymer physics have been borrowed and adapted to interpret experimental results.^3^ One such model is the “stickers and spacers model”, where different types of modules come to play different roles in the polymer solution.^7^ In the case of IDPs or IDRs, aromatic amino acid residues like Tyr or Phe behave as stickers that can form intra and intermolecular contacts, while glycine and polar amino acid residues are spacers that do not have strong interaction patterns. This simple model has been used successfully to interpret experiments on prion-like low complexity domains, which are frequently involved in the formation of condensates.^13^ Both experiments and simulations show that specific patterns of stickers and spacers in polypeptide sequences are important determinants of phase separation.^10,11,13–15^ Specifically, both randomly and regularly distributed stickers result in the formation of liquid-like condensates, while patchy regions with multiple stickers clustered together result in the formation of amorphous precipitates.^13^

The duality from the stickers and spacers model for different amino acid types may apply more broadly, even in the absence of an ordered polypeptide sequence where multivalency and patterning effects become important. In fact, composition alone may be sufficient to drive phase separation in the absence of additional information. Here we explore this possibility using atomistic molecular dynamics (MD) simulations. Molecular simulations with atomistic resolution of condensate formation have been attempted by others before.^16–18^ These simulations often involve multiple protein chains and large simulation boxes, which unfortunately cannot be run for sufficiently long timescales for observing phase separation or computing equilibrium properties. Alternatively, much more computationally cheap coarse-grained simulation models can recapitulate general principles of phase separation of disordered proteins,^9,19–23^ although details of chemical interactions are inevitably lost. An intermediate possibility is to use coarse-grained models with explicit solvent that may more faithfully account for hydration properties of condensates.^24–26^ Even in this case, protein-solvent interactions may need to be rebalanced and differences in the solvation entropy contribution need careful consideration due to reduction in the number of degrees of freedom from the coarse-graining. ^24^

In this work we run atomistic MD simulations using amino acid mixtures as a minimal, albeit detailed, model for macromolecular phase separation in protein-like systems. This type of simulations has been carried out in the past to study crowding and peptide solubility,^27–29^ although often in single component boxes. Because of the small size of amino acid residues, we can run direct coexistence simulations in solutions of hundreds of molecules. Specifically, we attempt to capture the behaviour that emerges from mixing amino acids of the sticker and spacer types, using compositions inspired by those observed in LCRs. Specifically, we choose Ser and Gly, which make over 50% of the sequence in some LCRs,^12^ as representative spacers, and the aromatic Tyr, which has been shown to be key for phase separation,^14^ as a sticker. From our simulations, we conclude that LLPS is an emergent property of amino acid mixtures driven by composition.

## Materials and Methods

### System Preparation

We have used the Packmol software^30^ to introduce multiple copies of acetylated and amidated amino acid residues, usually termed dipeptides, in elongated boxes with longest dimension (*L_x_*) between 10 and 20 nm, and *L_y_* = *L_z_* = 3.5 nm. We generate boxes for pure dipeptide solutions for Gly, Ser and Tyr residues, as well as stoichiometric binary and ternary mixtures. Target concentrations for these systems were between 1 and 2.5 M (see Supporting Information, Tables S1-S3). Amino acids were built using using the `leap` program from the Amber suite^31^ and then solvated to fill the simulation boxes. For most of our simulations, the Amber 99SB-ILDN force field^32^ together with the TIP3P water model^33^ was used for all systems. We have run additional simulations using two other force fields from the Amber family, ff03^⋆34^ and ff99SB-disp.^35^

### Molecular dynamics simulations

For equilibrium simulations, we followed the same simulation workflow irrespective of molecular system, box size or force field combination. The amino acid boxes were solvated, energy minimized and equilibrated in the NVT and NPT ensembles at 300 K. Equilibration involved an initial energy minimization step using a steepest descent algorithm, a short (500 ps) NVT simulation with position restraints on the protein heavy atoms to equilibrate the water molecules at 300 K and a 1 ns NPT run to converge the density to the corresponding value at 1 Pa using the Berendsen barostat.^36^ Long (500 ns-1 μs) production simulations were run for all systems at 300 K in the NVT ensemble. In all cases, a stochastic velocity-rescaling thermostat was used^37^ with a time step of 2 fs. Long range electrostatics were calculated using the particle-mesh Ewald method.^38^ We used the GROMACS program^39^ (version 2021) to run all the simulations.

For the boxes with *L_x_* =10 nm as longest dimension, we started our simulations from a random distribution of blocked amino acids. For longer boxes of *L_x_* = 20 nm, where a random mixture may easily result in the independent nucleation of condensates at multiple positions in the box, we run direct coexistence simulations, where a preformed condensate is first simulated in a shorter (*L_x_* = 8 nm long) box. Then the box was expanded to the final 20 nm length, filled with solvent molecules, and equilibrated in the NPT ensemble before the production run. In order to estimate dynamical properties of the resulting dense and dilute phases for selected systems, we sliced snapshots from the equilibrium runs and simulated them independently. In addition to the equilibrium runs, we have used the temperature replica exchange method to estimate the phase diagram of the Gly/Ser/Tyr equimolar mixture in the simulation box with 20 nm as longest dimension. Forty replicas with temperatures ranging between 288 and 444 K were initialized from the system equilibrated at 300 K and run in parallel for 300 ns with attempted replica swaps every 1000 integration time steps. The first 100 ns from each replica were discarded from the analysis.

### Analysis of the simulations

We have analyzed the simulation results using a combination of the suite of programs included as part of the GROMACS package^39^ and in-house Python scripts that make ample use of the MDtraj library.^40^ Molecular visualization was performed using the VMD program.^41^ Specifically, the solvent accessible surface area was calculated using the `gmx sasa` program.^42^ For the spontaneous phase separating systems, the resulting values were employed to estimate the *collapse degree*,^43^ defined as the ratio between the solvent accessible surface area in the initial configuration and that at a given time *t*. The number and distribution of molecular clusters were estimated using the `gmx clustsize` tool using a cutoff of 7 Å. The *clustering degree* was defined as the size of the largest cluster. For the collapse and clustering degrees we estimated errors from block averaging, leaving out the initial segment of the simulation where the collapse takes place.

We calculated density profiles across our simulation boxes using the `gmx density` program. Phase diagrams were determined from the temperature replica exchange data using the methods described by Dignon et al.^9,19^ For each temperature where we observe coexistence, we estimated the protein densities for the protein-dense and dilute phases, *ρ_H_* and *ρ_L_*, respectively. The critical temperature was then fitted from the equation

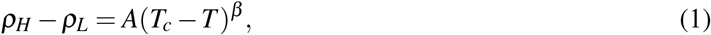

where *β* = 0.325 is the critical exponent^9,19^ and the critical density is derived from the law of rectilinear diameters

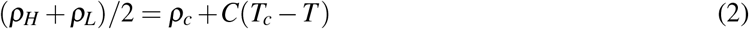

with positive fitting constant *C.*

Diffusion coefficients were calculated using the Einstein relationship, 〈|**r**(*t*+*τ*) – **r**(*t*)|^2^〉_*t*_= 6*Dτ*, where the brackets denote averages over different starting points, *t*. The fits of the diffusion coefficient we limited to the range of times to 1 to 5 ns where there are no deviations from linearity. Autocorrelation functions for the torsion angles were calculated from the simulation trajectories using the expression by Van der Spoel and Berendsen^44^

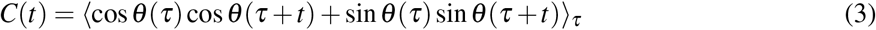

where *θ* is the angle of interest and *τ* is the lag time.

## Results and Discussion

### Composition determines phase behaviour in aminoacid solutions

First we focus on a set of simulations of pure amino acid solutions, where copies of the capped amino acid molecules were randomly inserted in 10 nm×3.5 nm×3.5 nm simulation boxes for residues of the spacer (Gly, Ser) and sticker (Tyr) types at concentrations between 1 and 2.5 M (see Fig. 1). In the case of the Gly boxes, we find that the peptide remains soluble even at the largest concentration for the full duration of the MD simulations, resulting in flat density profiles. For Ser, the dipeptide solution is mostly homogeneous up to 2 M, while at 2.5 M some regions become enriched in amino acid (Fig. 1, second column). Failing to form the condensate could be due to insufficient simulation time. For this reason we have run an additional simulation starting from a condensate. As monitored by the increase in the solvent accessible surface area SASA, the slab rapidly dissolves (see Fig. S1). On the contrary, in the case of Tyr, we find phase separation across all the concentrations explored, resulting in the formation of a slab (Fig. 1, center column). The differences observed between Gly/Ser and Tyr are consistent with the very low solubility reported for experiments on Tyr in its zwitterionic state^45^ and suggest that very small amounts of this amino acid may have drastic effects in properties of mixtures, as indeed has been found experimentally.^14^ We then repeat the same type of calculations but in this case for amino acid mixtures of spacers and stickers (see Fig. 1, fourth and fifth columns). We first look at equimolecular combinations of Gly and Ser for total amino acid concentrations between 1 and 2.5 M, as before (hence ranging between 0.5 and 1.25 M for each component). We find that these mixtures retain the properties of the single amino acid solutions of their constituents. At all concentrations probed in our calculations, the density profiles remain flat, which is consistent with the mixture exhibiting properties intermediate between their individual components. However, simulations for the same range of total concentration with stoichiometric amounts of Tyr result in the separation of the amino acid solution into dilute and dense phases.

**Figure 1:**
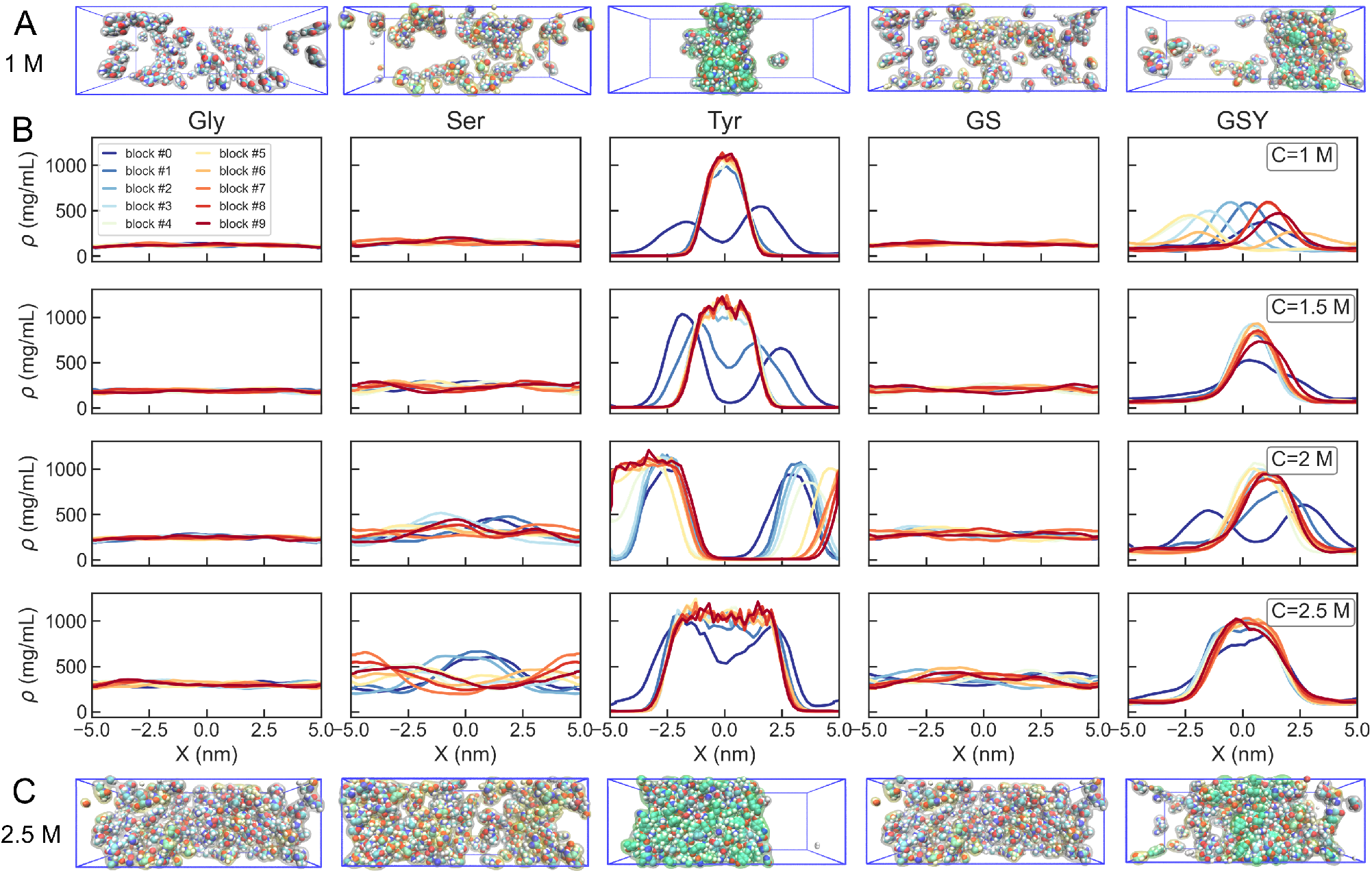
(A, C) Snapshots from equilibrium simulations at 1 M and 2.5 M. Left to right: Gly (white), Ser (yellow), Tyr (green) dipeptides, and their Gly/Ser and Gly/Ser/Tyr mixtures. simulations (first, second and third, respectively). (B) Density profiles along the longest axis of the simulation box for the pure dipeptide solutions at different concentrations for glycine (left column), serine (center) and tyrosine (right). Different colours correspond to averages over 50 ns blocks in the 500 ns simulations.

This result may be a consequence of the protein force field and water model combination used in our simulations. Although, the Amber ff99SB-ILDN force field with TIP3P water appropriately reproduces experimental solution densities in simulations of crowded amino acids,^27^ it also induces compact states in proteins.^46^ We have run simulations with two other optimized force fields of the Amber family at 2 M amino acid concentration (see Figure S2 in the Supporting Information). Using the ff03^⋆^ force field^34^ with TIP3P water, the density profiles for the Gly/Ser/Tyr mixture are comparable to those obtained using ff99SB-ILDN, indicating that our result applies beyond our chosen force field and water model combination. With the more recently derived a99SB-disp force field and water,^32^ the amino acid mixture fails to phase-separate. The latter result can be attributed to the same reasons that prevent this specific force field and water model combination to adequately capture protein-protein binding, which led to its reparametrization. ^47^ In any case, the properties of the simulated amino acid condensates will depend on the choice of the force field.

### Clustering degree and aggregation propensity as metrics of condensate formation

In all systems for which we report the formation of condensates (i.e. Tyr and Gly/Ser/Tyr), the initial collapse takes place within the first 50-100 ns of the simulation run. After this initial collapse, the systems sample the equilibrium distribution for the remainder of the 500 ns runs, suggesting good convergence in our simulations (see Supplementary Fig. S3). In Figure 2A, we show the time series for the solvent accessible surface area (SASA) for all systems at 1.5 M concentration. In the case of the Gly/Ser/Tyr mixture, *SASA* decreases to a value intermediate between that of the Gly/Ser solutions and that of the Tyr simulation box. This result occurs consistently across the full range of concentrations explored in our simulations (see Supplementary Fig. S3). This type of intermediate behaviour for the Gly/Ser/Tyr mixture is also observed for the maximum cluster size, which has also been shown to be a robust descriptor of the degree of clustering^26^ (Fig. 2A). We note that also the time series data for the maximum cluster size and the average number of clusters seem stable after the initial 100 ns of simulation, which gives further reassurance on the convergence of the results (see Supplementary Figs. S4 and S5). The only exception is the 2 M solution of Tyr, which nucleates in two different points along the simulation box, forming two slabs that finally merge at ~250 ns.

**Figure 2:**
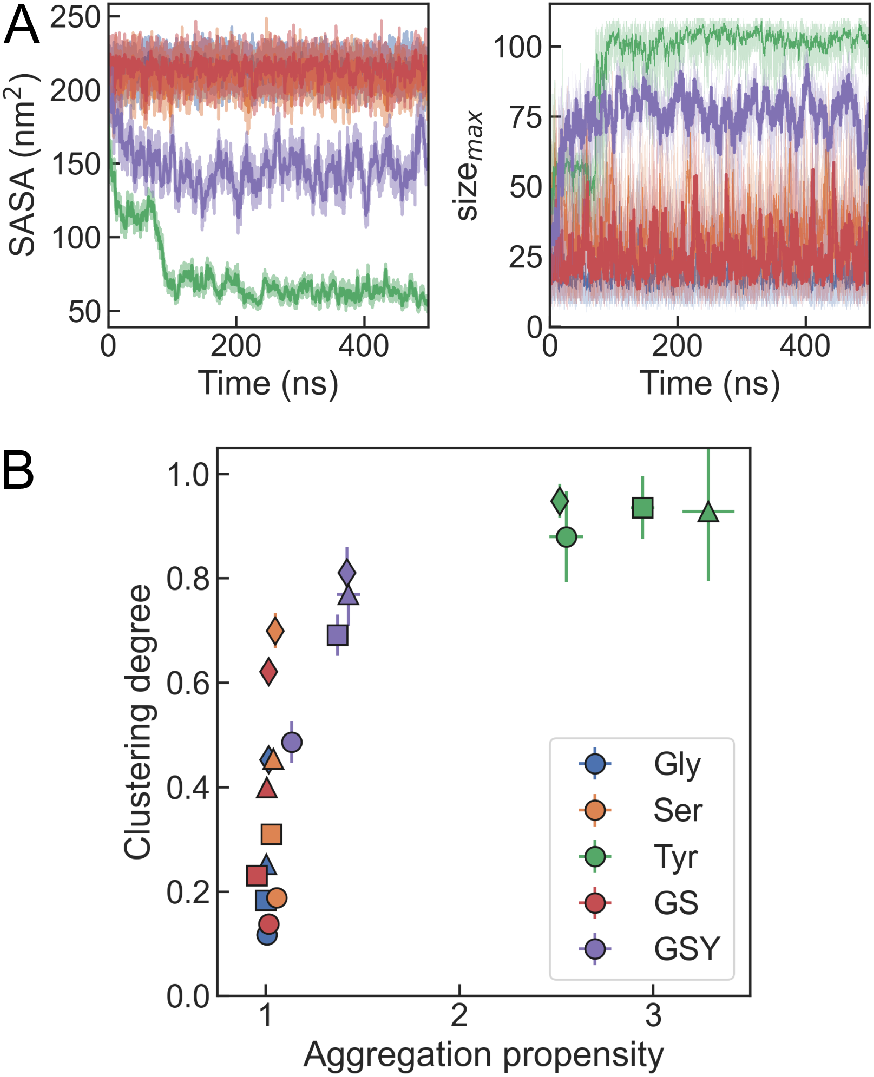
(A) Time series data for the solvent accessible surface area (*SASA*, top) and the clustering degree (bottom) for all systems in the 1.5 M solutions. (B) Projection of all the simulation datasets on the aggregation propensity and clustering degree. Different symbols total peptide concentrations (circles: 1 M; squares: 1.5 M; triangles: 2 M; diamonds: 2.5 M).

Two useful metrics that summarize the information from these simulations are the “aggregation propensity”, defined as the ratio between the initial and instantaneous solvent accessible surface areas,^43^ and the “clustering degree”, which we obtain normalizing the maximum cluster size by the total number of residues in the box.^26^ In the projection of our datasets on these axes (Fig. 2B), the Gly/Ser/Tyr system stands out as having intermediate values of the aggregation propensity and clustering degree between the systems that do not phase separate (Gly, Ser and Gly/Ser mixture) and the Tyr boxes. This result suggests a qualitative different behaviour between the sticker-only or spacer-only solutions and the mixture of stickers with spacers.

### Densities from coexistence simulations

Despite the interesting differences identified from these simulation runs, the relatively small size of our simulation boxes may lead to finite size effects. This is noticeable, particularly, for the lowest concentration (1 M). As a result, for the Gly/Ser/Tyr mixture, the density peaks at different values depending on the target concentration. In order to address the severity of these problems we have run additional simulations for longer simulation boxes with longest dimension of *L_x_* = 14 nm. For the 2 M boxes of all single and multi-component systems, we find consistent results irrespective of box size (see Supplementary Fig. S6). For the multi-component Gly/Ser/Tyr system, which we are most interested in, we have explored the dependence with the box length across concentrations. For even longer boxes we often find simultaneous nucleation of condensates at multiple points along the X-axis that fail to merge within the simulation time.

In order to accurately determine the densities of amino acid condensates, we resort to the slab method where we start from a pre-equilibrated biomolecule-dense phase that is put in contact with bulk water (see Methods). Specifically, we run simulations using the temperature-replica exchange method for long boxes of *L_x_* =20 nm at 2 M total amino acid concentrations, from which we calculate density profiles (see Figure 3). We observe the expected temperature dependence for both protein and water densities, with gradually disappearing slabs upon increasing temperature. The results are consistent with those presented above for the three component mixtures in the shorter boxes. At room temperature, the densities of the protein-dilute and protein-dense phases are, respectively, ~50 mg/mL and ~ 1040 mg/mL. Conversely, the water density in the condensate is only ~ 180 mg/mL. These values for the densities are more extreme than those observed for the FUS low complexity domain and LAF-1 RGG protein in experiments and previous atomistic simulations, where condensates where typically much more hydrated (e.g. having ~500 mg/mL protein and ~600 mg/mL water in the dense phase^17^). This difference may be due to the tighter packing that becomes possible in our systems, due to the reduction in excluded volume effects from a covalently bonded polypeptide chain, and may also be influenced by the combination of force field and water model.

**Figure 3:**
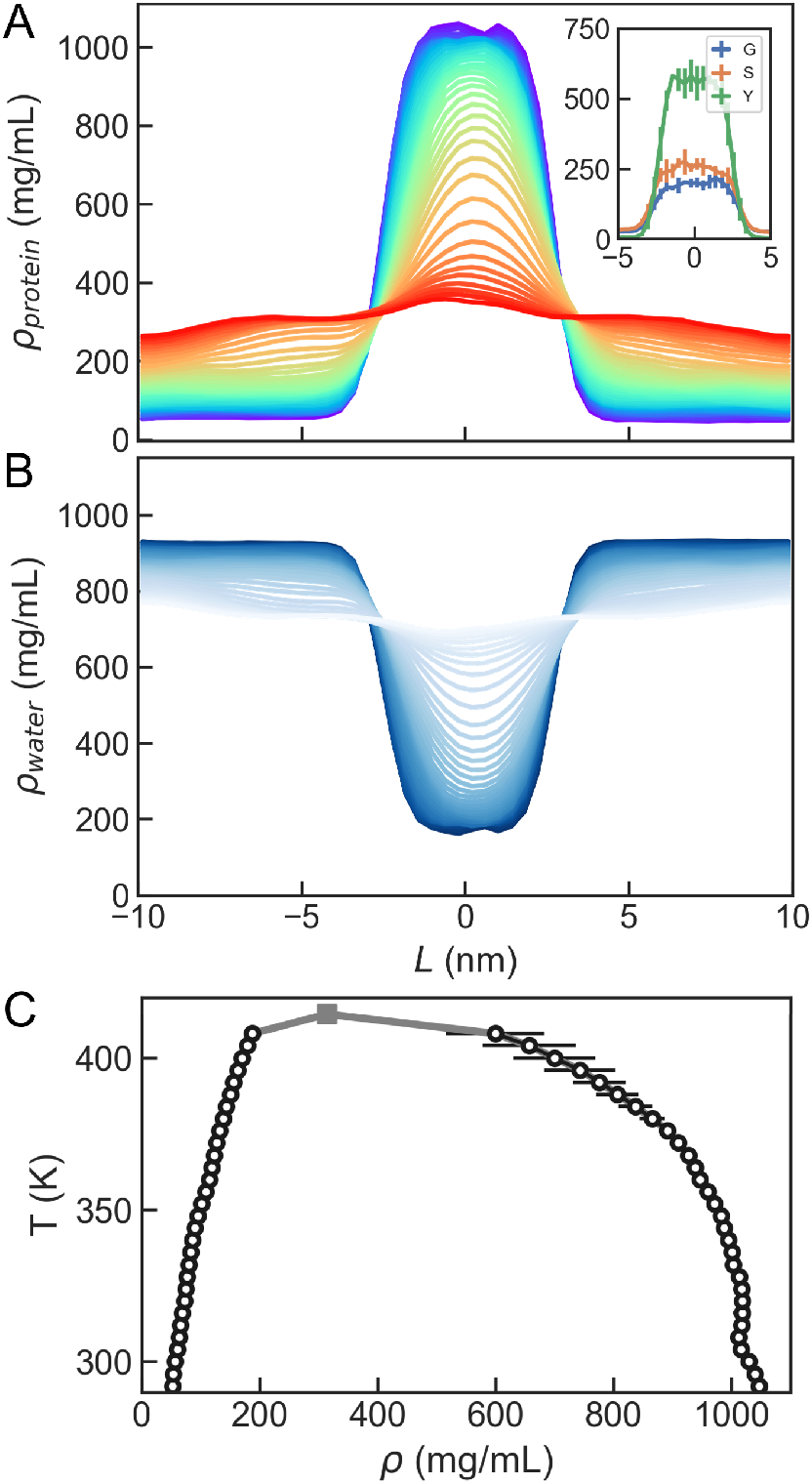
(A) Density profiles from replica exchange MD simulations of the ternary Gly/Ser/Tyr equimolecular mixture at 2 M. Blue and red lines corresponds to the lowest and highest temperatures, respectively. Inset: density profiles for Gly, Ser and Tyr at room temperature. (B) Density profiles for water molecules for all replicas. Lighter colours correspond to higher temperatures. (C) Phase diagram derived from the REMD simulations (circles). The fitted critical point is shown as a square.

### Phase behaviour in the ternary amino acid mixtures

We have seen that our results from simulations of very simple mixtures of capped “sticker and spacer” amino acids captured properties observed in many disordered protein segments, which experience a phase transition with an upper critical solution temperature (UCST).^12^ This type of phase behaviour is recapitulated in our simulations. We use the densities of the dense and dilute phases (*ρ_H_* and *ρ_L_*, respectively), from the temperature dependent density profiles obtained from the replica exchange simulations and derive a phase diagram that can be fitted to extract a critical temperature of *T_c_* = 414 K (see Fig. 3C).

So far we have treated the amino acid mixtures as homogeneous protein solutions. However, lacking the connectivity of covalently-bonded polypeptide chains, amino acids could be partitioning independently of each other. We have used the density profiles for Gly, Ser and Tyr to calculate their individual compositions along the simulation boxes (see Fig. 3A, inset). Although all three amino acids are enriched in the slab, the condensed phase is richer in Tyr than in Gly/Ser, which is consistent with the the greater hydrophobicity of this amino acid. Conversely, the dilute phase is richer in Gly/Ser while the Tyr density of this phase is almost negligible.

In order to map the composition dependence of the phase behaviour, we have run 1 *μ*s equilibrium simulations for amino acid mixtures varying the ratio between stickers and spacers while retaining the total number of amino acid residues in the mixtures constant. From the densities of sticker and spacer types (*ρ_Y_* and *ρ_GS_*, respectively) in the dilute and dense phases, we obtain the phase diagram shown in Figure 4A (we also show representative snapshots from simulations at different compositions in Figure 4B). The emerging picture is that of a “scaffold-client” system,^48^ where the spacer-type residues, which do not phase-separate on their own, are recruited by the stickers. This is due to the greater strength of the homotypic interactions between tyrosine residues, relative to every other pairwise interactions. However, heterotypic interactions between tyrosine and glycine/serine also contribute to the stability of the condensate. This is supported by the statistics of pairwise contacts among residues (which we show for the equimolar mixture in Fig. S7 of the Supporting Information). While Tyr is the residue that forms more contacts, these are not exclusively homotypic, as Tyr-Ser and Tyr-Gly contacts are also relevant. We note that the relative weights of different types of contacts is comparable to that observed by Zheng and his co-workers in atomistic simulations of FUS LC and LAF1 RGG (i.e. with Tyrosine contributing twice as much as other residue types). ^17^ Our results support that spacers are not inert but relevant actors in the phase-behaviour of protein condensates.

**Figure 4:**
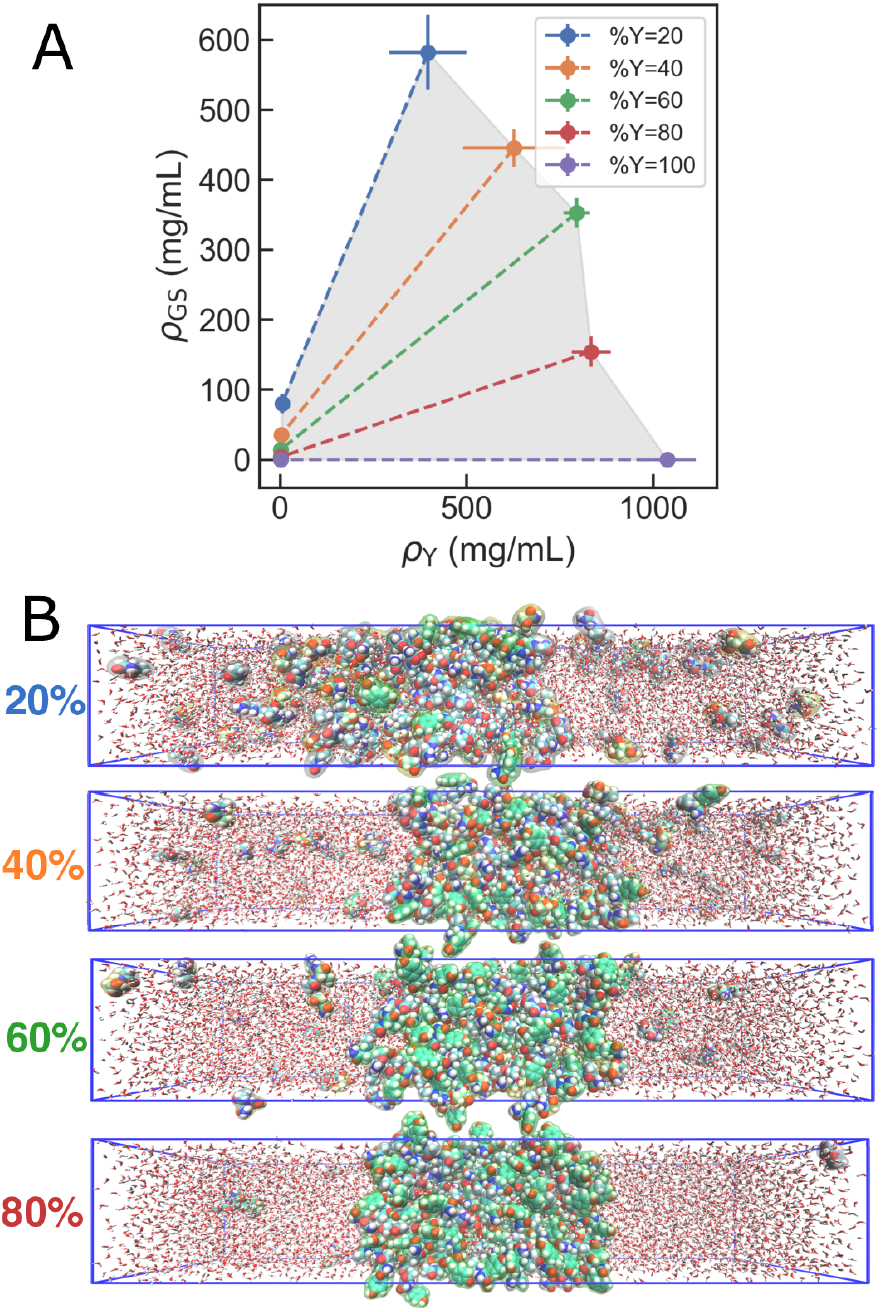
(A) Two-component phase diagram from simulations of Gly/Ser/Tyr mixtures for different compositions, represented in terms of sticker and spacer densities (*ρ*_Y_ and *ρ*_GS_, respectively) for the dilute and dense phases. Different colours indicate simulated fraction percentage of Tyr within the mixture. (B) Representative snapshots of the simulation for different compositions.

### Slowdown in the dynamics within the molecular condensates

An important consideration is to which extent the condensates formed are aggregates or retain a degree of fluidity comparable to the amino acids in solution. Even if a neat distinction may be difficult to establish, we can inspect the dynamical properties in our simulations to clarify this point. Calculating these from the coexistence runs would result in averaging their values across different phases. For this reason, we have run separate simulations of the condensed phases derived from pure Tyr and Gly/Ser/Tyr slabs and from the dilute phase of the ternary mixture. First, we use Einstein’s relation to calculate diffusion constants, *D*, for the amino acid residues in the condensed phases (see Fig. 5A). We observe that the value of *D* is considerably slower in the case of the Tyr condensate than in the Gly/Ser/Tyr mixture, with individual amino acid values commensurate with their size and stickiness. This result indicates a greater slowdown in the dynamics in the Tyr condensate than in the ternary mixture. We have also calculated *D* for the water molecules in the condensates, which exhibit a drastic slowdown with respect to *D* for water in the dilute phase (4.4 nm^2^/ns) (see Fig. 5B). This effect is again more pronounced in the case of the Tyr condensate (3.7e-2 nm^2^/ns) than in the ternary mixture (6.7e-2 nm^2^/ns). These values are comparable to the slowest mode identified by Zheng and co-workers and attributed to water molecules interacting with protein atoms,^17^ which is an accurate description for that water molecules in our protein-rich condensates. From these simulations we also analyze the tyrosine dynamics by looking at the autocorrelation functions of the Ramachandran angles (*ϕ* and *ψ*) and the *χ*_1_ sidechain torsion (see Fig. 5C-E). When compared with the dilute phase the dynamics in the condensates are again slowed down, but to a larger degree in the Tyr condensates than in the ternary mixtures.

**Figure 5:**
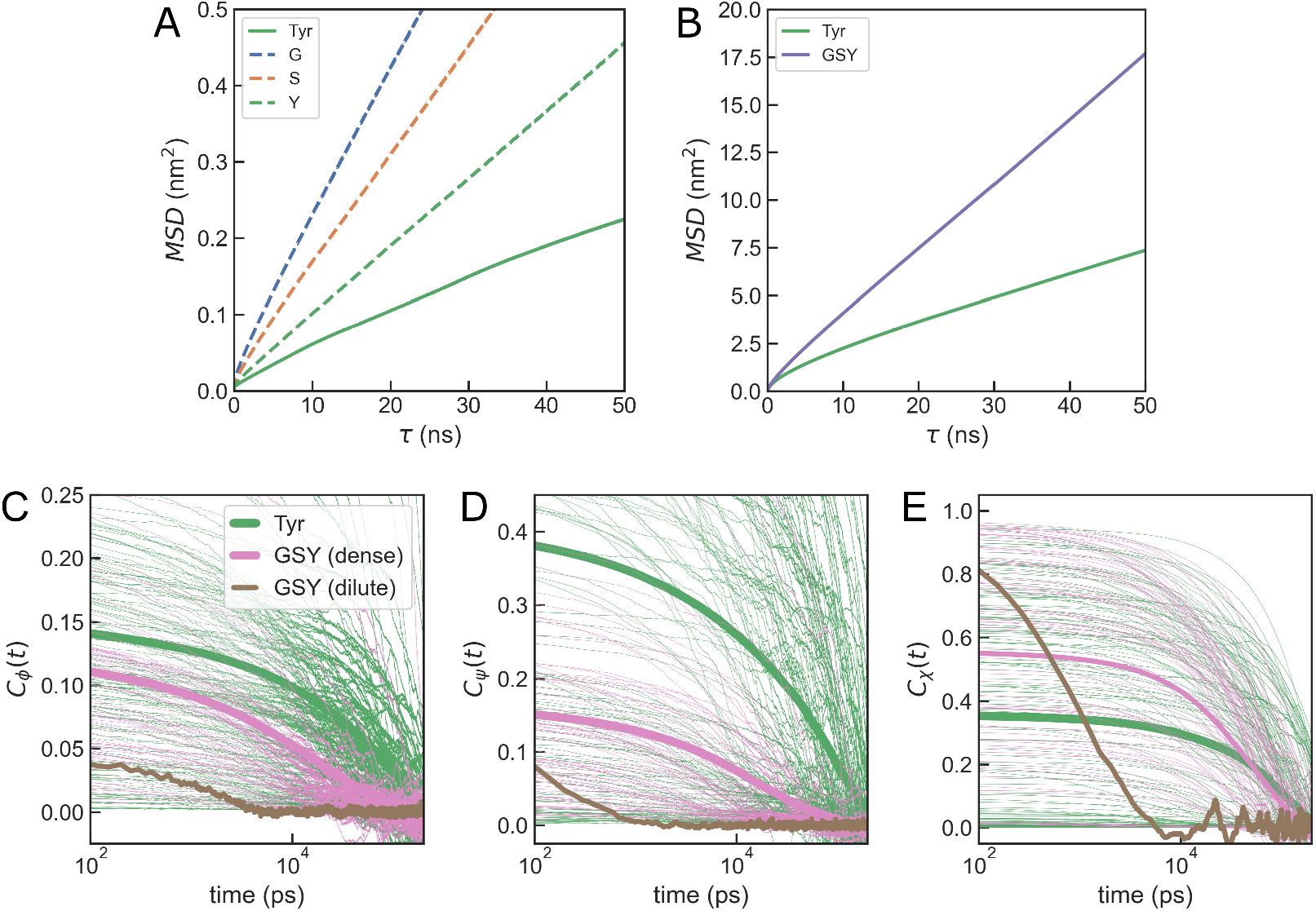
(A) Mean squared displacement vs lag time for amino acid residues in condensate simulations. The solid line corresponds to a Tyr condensate while dashed lines correspond to different components in the Gly/Ser/Tyr condensate. (B) Same for water molecules within the Tyr (green) and Gly/Ser/Tyr (purple) condensates. (C-D) Autocorrelation functions for the Ramachandran angles (*ϕ*, *ψ*) and first sidechain torsion for Tyr residues in the Tyr condensate simulations (green) and in the Gly/Ser/Tyr simulations for the dense (pink) and dilute (brown) phases. Thin lines are autocorrelation functions from individual molecules and thick lines are their averages.

### Conclusions

In summary, in this work we have showed with a minimalistic model system that LLPS is an emergent property of amino-acid mixtures that is governed by composition. Considering a mixture of “spacer” residues (Gly/Ser) as a reference system –which is warranted by the abundance of these amino acids in phase-separating LCRs– we show that small amounts of a “sticker” (in this case the rather insoluble Tyr amino acid) make their solutions separate into two phases with liquid-like properties. This suggests that, the duality of amino acid types in the “sticker and spacers” model holds more generally than in the context of IDPs and that the stoichiometry of the mixture alone may be predictive of the phase behaviour. Provided a small amount of aromatics or “stickers” is present, amino-acids, peptides or proteins may be able to spontaneously phase separate. The critical role of Tyr in our simulations is consistent with previous experiments on folded modules and intrinsically disordered regions. ^11,13,14^

The formation of aggregates as opposed to liquid-like condensates in the case of patchy sequences with clusters of stickers found experimentally^13^ may be understood as being closer to the results from our simulations of Tyr-only boxes. In this case, we observe the formation of condensates with greater degree of dynamical arrest, as monitored by both diffusion constants and intramolecular dynamics. The number of contacts formed by stickers remains stable after the formation of condensates, suggesting that ageing or maturation are not taking place at least in the timescales that we investigate.^25,49^

Our work also confirms in full atomistic detail recent results by Tang et al,^26^ who explored dipeptide solutions in simulations using the intermediate-resolution MARTINI model.^50^ Further evidence for the generality of LLPS is accumulating in the experimental front in a broad range of length-scales, from short peptides^51,52^ to globular proteins.^8^ The occurrence of macromolecular phase separation may hence turn out to be a general property of both peptides and proteins, determined by composition and further fine-tuned by multivalency and patterning,^13^ the presence of charged residues,^11^ nucleic acids,^53,54^ ionic strength^22^ or crowding agents.^55^

## Acknowledgement

Financial support to DDS comes from Eusko Jaurlaritza (Basque Government) through the project IT1254-19, Grants RYC-2016-19590 and PID2021-127907NB-I00 from the Spanish Ministry of Science and Universities through the Office of Science Research (MINECO/FEDER) and the Donostia International Physics Center (DIPC). The author thanks Xabier López for useful discussions and Athi N. Naganathan and Robert B. Best for their comments on the manuscript. The author also acknowledges the staff at the DIPC Supercomputing Center for technical support.

