## Supporting Information for "Phase separation in amino acid mixtures is governed by composition"

### Supporting Information for Liquid-liquid phase separation in amino acid mixtures is governed by composition

David De Sancho*

*Polimero eta Material Aurreratuak: Fisika, Kimika eta Teknologia, Kimika Fakultatea, UPV/EHU & Donostia International Physics Center (DIPC), PK 1072, 20018 Donostia-San Sebastian, Euskadi, Spain*

### List of Figures

S1 Time series data for the accessible surface area of the Ser 2.5 M solution started from a random distribution of blocked serine peptides (blue) and from a slab configuration (orange). 4

S2 Density profiles for three different force fields for the Gly/Ser/Tyr equimolar mixture. Results are shown for 10 blocks, 50 ns each, from the equilibriumn simulation trajectories, starting in lighter and ending in darker shades of the corresponding colour. . . . . . . . . . 4

S3 Time series data for the solvent accessible surface area (*SASA*) for all systems in the 1-2.5 M amino acid solutions. Running averages are shown in darker shades of colour. . . . . . . 5

S4 Time series data for the maximum cluster size for the Gly/Ser/Tyr equimolar systems at 1-2.5 M total amino acid concentrations. . . . . . . . . . . . . . . . . . . . . . . . . . . . 6

S5 Time series data for the average cluster size for the Gly/Ser/Tyr equimolar systems at 1-2.5 M total amino acid concentrations. . . . . . . . . . . . . . . . . . . . . . . . . . . . . . 7

S6 Density profiles for all systems at 2 M concentrations for simulation boxes of different length for the longest dimension. Each line corresponds to averages over 50 ns blocks. Darker lines correspond to longer times in the simulation. . . . . . . . . . . . . . . . . . . . . . . . . . . 8

S7 Average number of pairwise contacts for different residue types at 2 M peptide concentration for the GSY replica exchange simulations. A contact is defined whenever any pair of heavy atoms is closer than 6 Å. Bar charts correspond to the number of contacts per residue for each amino acid type. . . . . . . . . . . . . . . . . . . . . . . . . . . . . . . . . . . . . . 9

Table S1: Number of copies of each amino acid type for the different final concentrations in the case of $L_x = 10$ nm boxes of pure terminally blocked amino acids, and their binary and ternary mixtures.

| Total concentration (M) | Pure | Binary | Ternary |
|---|---|---|---|
| 1 | 73 | 36 | 24 |
| 1.5 | 110 | 55 | 36 |
| 2 | 147 | 73 | 49 |
| 2.5 | 184 | 92 | 61 |

Table S2: Number of copies of each amino acid type for the different final concentrations in the case of pure terminally blocked amino acid solutions, and binary and ternary mixtures for $L_x = 14$ nm.

| Total concentration (M) | Pure | Binary | Ternary |
|---|---|---|---|
| 2 | 206 | 103 | 68 |

Table S3: Number of copies of each amino acid type for the different final concentrations for the $L_x = 20$ nm boxes for the ternary mixtures.

| Total concentration (M) | Number of residues |
|---|---|
| 2 | 98 |

Table S4: Number of copies of each amino acid type for the different final concentrations for the $L_x = 20$ nm boxes for the ternary mixtures with different proportion of Tyr.

| Fraction Tyr | Number of Gly/Ser | Number of Tyr |
|---|---|---|
| 0.2 | 118 | 59 |
| 0.4 | 88 | 118 |
| 0.6 | 59 | 177 |
| 0.8 | 29 | 236 |
| 1 | 0 | 295 |

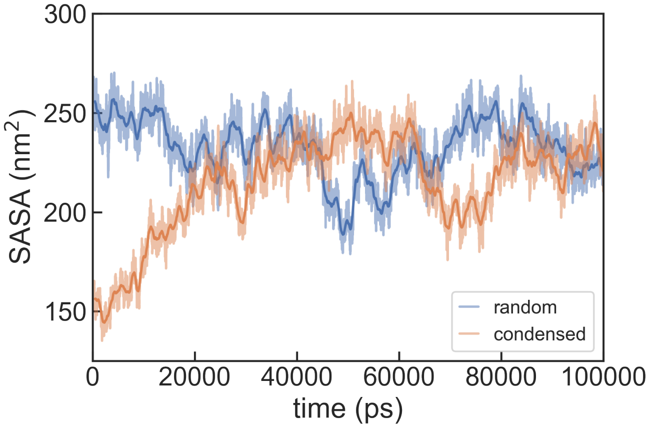

Figure S1: Time series data for the accessible surface area of the Ser 2.5 M solution started from a random distribution of blocked serine peptides (blue) and from a slab configuration (orange).

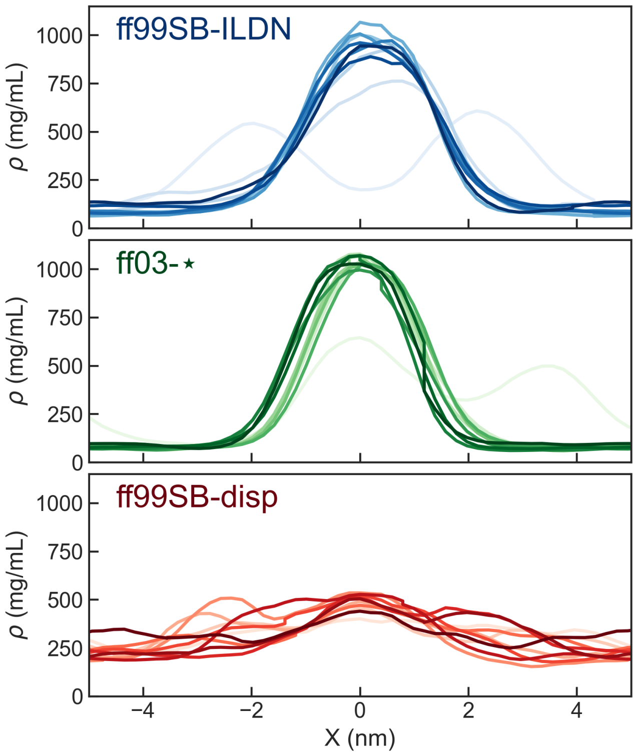

Figure S2: Density profiles for three different force fields for the Gly/Ser/Tyr equimolar mixture. Results are shown for 10 blocks, 50 ns each, from the equilibriumn simulation trajectories, starting in lighter and ending in darker shades of the corresponding colour.

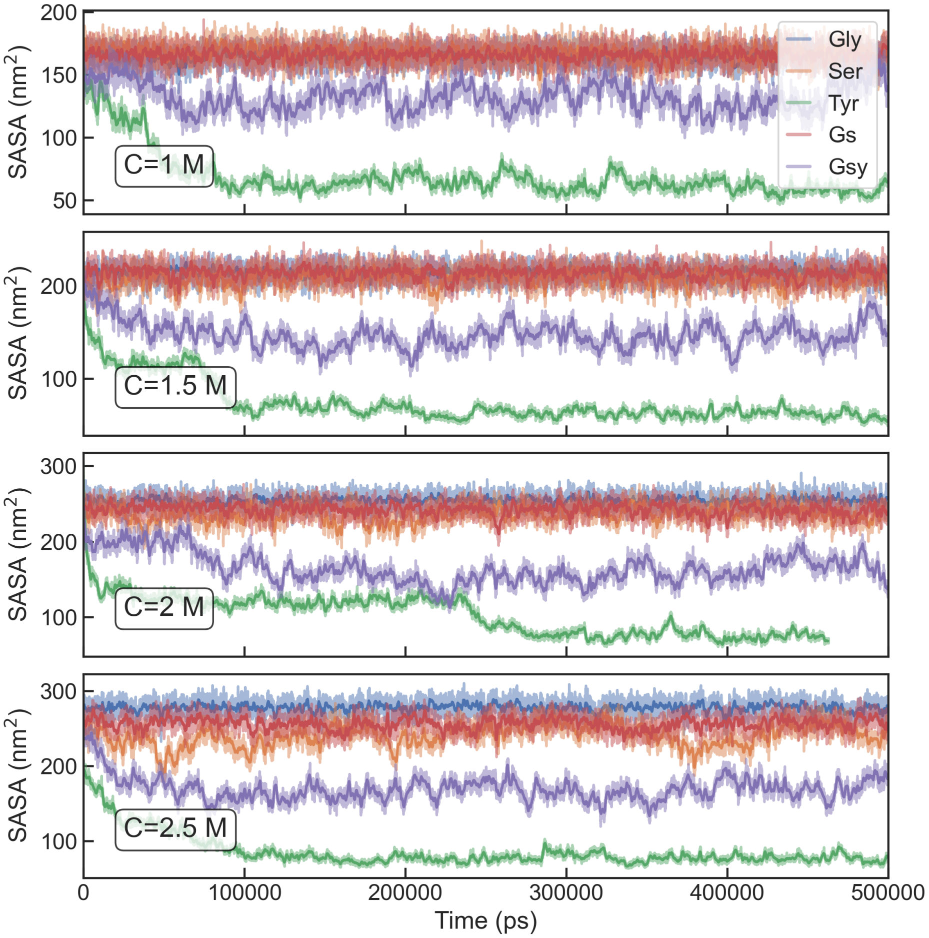

Figure S3: Time series data for the solvent accessible surface area (*SASA*) for all systems in the 1-2.5 M amino acid solutions. Running averages are shown in darker shades of colour.

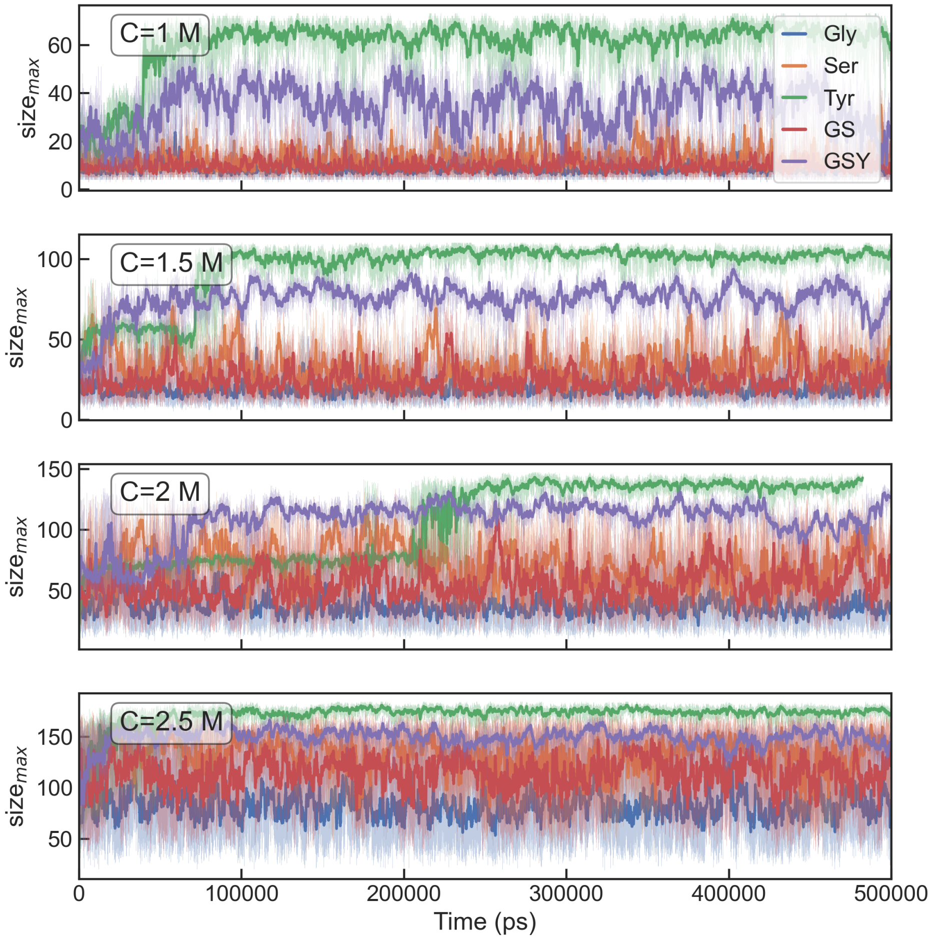

Figure S4: Time series data for the maximum cluster size for the Gly/Ser/Tyr equimolar systems at 1-2.5 M total amino acid concentrations.

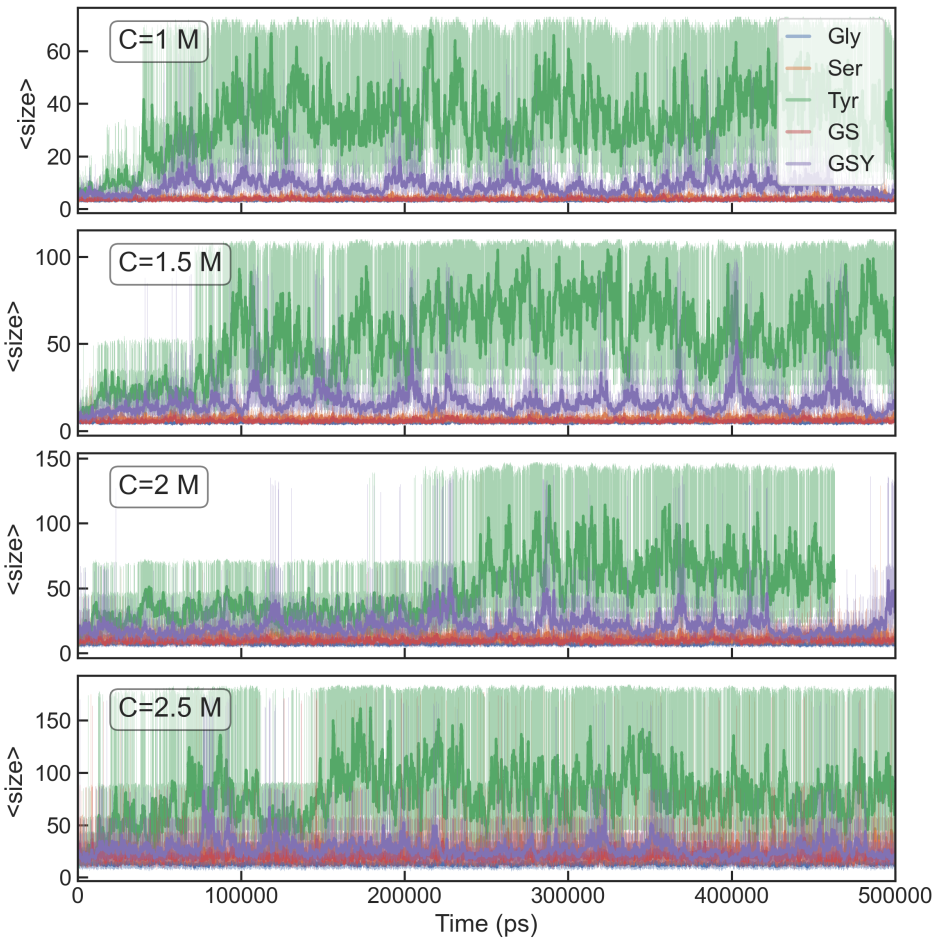

Figure S5: Time series data for the average cluster size for the Gly/Ser/Tyr equimolar systems at 1-2.5 M total amino acid concentrations.

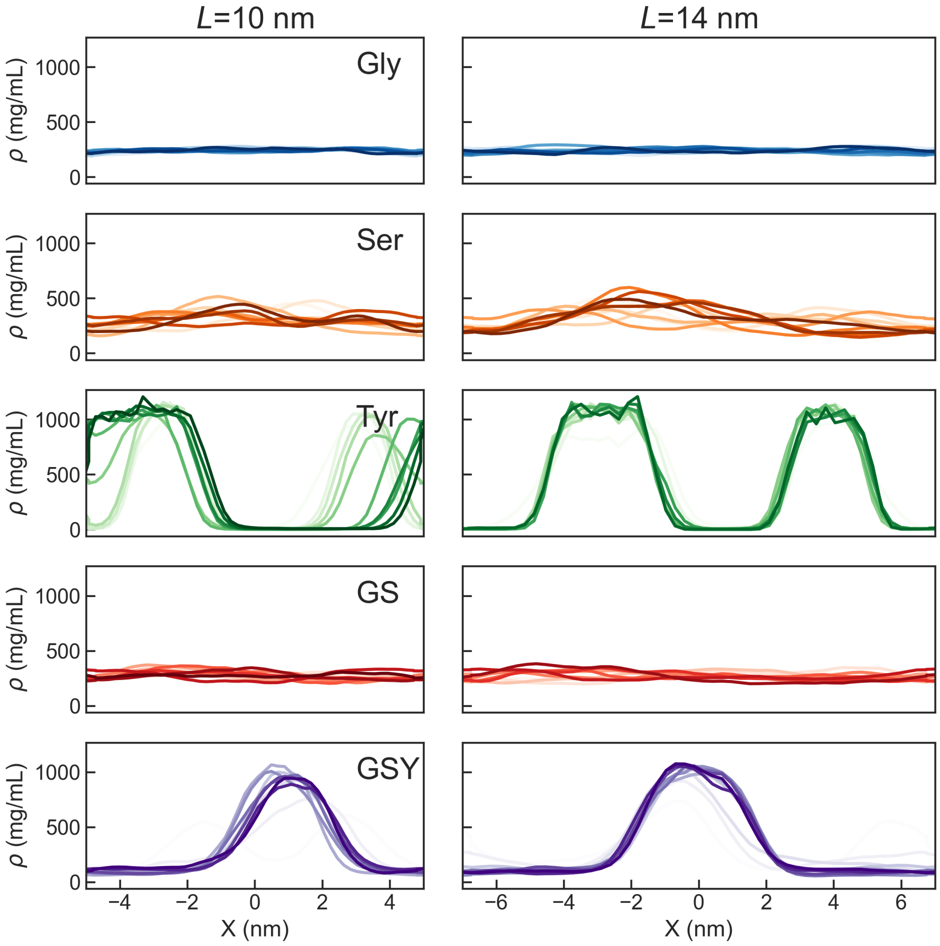

Figure S6: Density profiles for all systems at 2 M concentrations for simulation boxes of different length for the longest dimension. Each line corresponds to averages over 50 ns blocks. Darker lines correspond to longer times in the simulation.

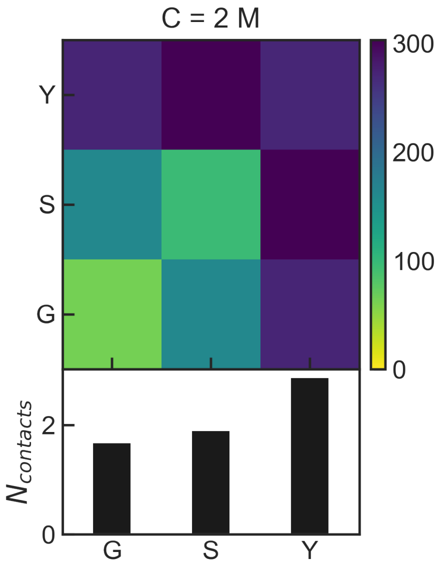

Figure S7: Average number of pairwise contacts for different residue types at 2 M peptide concentration for the GSY replica exchange simulations. A contact is defined whenever any pair of heavy atoms is closer than 6 Å. Bar charts correspond to the number of contacts per residue for each amino acid type.
